# Phe-Gly motifs drive fibrillization of TDP-43’s prion-like domain droplets

**DOI:** 10.1101/2020.02.20.957969

**Authors:** David Pantoja-Uceda, Cristiana Stuani, Douglas V. Laurents, Ann E. McDermott, Emanuele Buratti, Miguel Mompeán

**Affiliations:** “Rocasolano” Institute for Physical Chemistry, Spanish National Research Council, Serrano 119, 28006, Madrid, Spain; International Centre for Genetic Engineering and Biotechnology (ICGEB), Padriciano 99, I-34149, Trieste, Italy; Department of Chemistry, Columbia University, New York, NY 10027, USA

**Author notes:** These authors contributed equally to this work (David Pantoja-Uceda, Cristiana Stuani).

## Abstract

TDP-43 assembles various aggregate forms, including biomolecular condensates or functional and pathological amyloids, with roles in disparate scenarios (*e.g*. muscle regeneration versus neurodegeneration). The link between condensates and fibrils remains unclear, just as the factors controlling conformational transitions within these aggregate species: salt- or RNA-induced droplets may evolve into fibrils or remain in the droplet form, suggesting distinct endpoint species of different aggregation pathways. Using microscopy and NMR methods, we unexpectedly observed *in vitro* droplet formation in the absence of salts or RNAs, and provided visual evidence for fibrillization at the droplet surface/solvent interface but not the droplet interior. Our NMR analyses unambiguously uncovered a distinct amyloid conformation in which Phe-Gly motifs are key elements of the reconstituted fibril form, suggesting a pivotal role for these residues in creating the fibril core. This contrasts the minor participation of Phe-Gly motifs in initiation of the droplet form. Our results point to an intrinsic (*i.e*., non-induced) aggregation pathway that may exist over a broad range of conditions, and illustrate structural features that distinguishes between aggregate forms.

---

Transactive response DNA-binding Protein of 43 kDa (TDP-43) is an RNA-binding protein that forms aberrant aggregates associated to disease contexts, including frontotemporal dementia (FTD) and amyotrophic lateral sclerosis (ALS)^1^, or the recently reported “LATE” dementia, which had been misdiagnosed as Alzheimer’s disease^2^. TDP-43 also aggregates into functional amyloids^3^, and participates in the assembly of various biomolecular condensates^4,5^, which highlight the need for structural studies to understand the interplay between these aggregate forms. Structurally, TDP-43 contains a well-folded N-terminal^6^ and two RRM domains^7^, and a low complexity, prion-like domain (PrLD) at its C-terminus, which seems pivotal for the formation of the diverse amyloid and droplet forms. The TDP-43 PrLD is large (residues 267-414), comprising almost half the length of TDP-43, and is intrinsically disordered, with the exception of a short hydrophobic segment (residues 318-340) that contains a helical region. This hydrophobic helix is followed by a Q/N-rich stretch (residues 341-367), and all this 318-367 (hydrophobic+Q/N) stretch is N- and C-terminally flanked by two segments rich in Gly, Aromatic, and Ser residues (“GAroS” segments, residues 273-317 and 368-414)^8^. Consensus exists that TDP-43 PrLD droplets or condensates can be induced with NaCl or RNA molecules, and that structurally, droplet assembly is orchestrated by the hydrophobic helix^9-11^, with moderate assistance of Trp and other aromatic residues within the two GAroS segments^11^. Importantly, this prior, seminal work did not detect maturation of the droplets into amyloid-like fibrils^9-12^. Conversely, Lim *et al*. reported that TDP-43 PrLD assembles amyloid fibrils, also driven by the hydrophobic helix, albeit they did not detect droplet formation^13^. These important observations suggest that droplet and fibril formation are end points of two distinct aggregation pathways, which intriguingly involve the same region within TDP-43 PrLD; namely, the hydrophobic helix. However, this dichotomous picture (*i.e*., droplets *or* fibrils) contrasts with recent findings that proved that the droplet environment is conducive for fibril formation^14,15^, as broadly established for other PrLD-containing proteins with GAroS regions^16-18^.

Using polypeptide constructs corresponding to the TDP-43 PrLD central region (residues 311-360) that contains the critical helix, but excludes the two flanking GAroS segments (residues 267-310 and 361-414), the Eisenberg lab obtained cryo-EM structures of polymorphic fibrils with a common long β-hairpin named a “dagger-shaped” fold^19^. A most recent solid-state NMR (SSNMR) study on fibrils from this same 311-360 construct revealed key intermolecular contacts that are not compatible with the long β-hairpin structure seen by cryo-EM^15^, which supports a distinct fibril conformation. Interestingly, these fibrils stem from droplets^15^. Although these constructs correspond to a limited part of the TDP-43 PrLD sequence as they exclude the two GAroS segments, these model fibrils already reveal the rich diversity of the TDP-43 PrLD amyloid foldome, which assembles distinct functional (*e.g*. in muscle regeneration)^3^ and pathological (*e.g*. in FTD)^20,21^ amyloid-like structures. However, we do not know if the structures formed by the TDP-43 PrLD (311-360) constructs recapitulate those adopted by the entire TDP-43 PrLD, which contains the two flanking GAroS regions (*i.e*., 267-310 and 361-414) that are relevant for droplet formation^11^.

All the above observations pertinent to TDP-43 PrLD aggregation pathways have been carried out at pH values ranging from 6.0 to 8.0, and rely on droplet induction by salt or RNA molecules^9-12,14^. These efforts have lead to the successful elucidation by NMR of the helix-driven, GAroS-assisted mechanism of droplet assembly at the residue-level^9-12^. However, we lack such fundamental knowledge on the *intrinsic* aggregation (*i.e*., in the absence of inducers) of TDP-43 PrLD at low pH. So far, only Lim *et al*. have reported an NMR characterization of the TDP-43 PrLD at pH 4, and they found no aggregation at all within months, even at protein concentrations as high as 600 µM^13^. This is in contrast with the observations we inform on this manuscript, where TDP-43 PrLD aggregates at pH 4, even when no salts or other aggregation inducers (*e.g*. RNAs) are present. Intense metabolic activity lowers pH, which induces stress granule formation^22-24^ where TDP-43 is reported to form harmful ALS-relevant aggregates^5,25^. These aspects of TDP-43 aggregation in pro-pathological and pro-physiological contexts prompted us to structurally characterize the aggregation pathway of TDP-43 PrLD at pH 4 at the residue level, using liquid state (LS) and solid-state (SS) NMR spectroscopies.

In the following, we report three fundamental aspects of TDP-43 PrLD; namely, (i) that the full TDP-43 PrLD (residues 267-414) form droplets at pH 4 and in the absence of ions or RNA molecules, (ii) that these droplets do not represent endpoint species from this intrinsic aggregation pathway, but instead afford amyloid accumulation at the droplet surface/solvent interface, and (iii) that such amyloid cores are stabilized by Phe/Tyr, Gly, and Ser residues from GAroS regions, located outside the central region containing the hydrophobic helix.

## Results

### The TDP-43 PrLD intrinsically self-aggregates at pH 4 to enable droplet formation

We observed that the PrLD of TDP-43 aggregated at pH 4 in the absence of salt or RNA, which, to the best of our knowledge, was unexpected^13,14^. Because induced aggregation at higher pH values is reported to be driven by the central region containing a helical segment^9,10^, we sought to confirm whether this general mechanism is also operative under our conditions, using liquid state NMR (LSNMR).

Considering that the TDP-43 PrLD is of low complexity (*i.e*. poor amino acid variability), aggregation-prone, and chiefly disordered, the LSNMR ^13^CO, ^13^Cα, ^13^Cβ, ^15^N, ^1^HN, ^1^Hα and ^1^Hβ correlations were obtained using a non-conventional strategy that affords robust and complete assignments with a minimal set of experiments. In brief, this approach is based on the unambiguous determination of sequential connectivities between all spin systems (except prolines) by direct correlation of the ^1^HN and ^15^N amide groups of one amino acid residue to those of the next residue in the sequence, followed by the assignment of each ^13^Cα and ^13^Cβ nuclei to corroborate the identity of the sequential fragments interrupted by prolines (see *Methods* and **Table S1** and **S2** for a more detailed description). This methodology proves useful for low complexity stretches and aggregation-prone samples, and allowed us to assign all residues under the conditions of this study (BMRB entry 50154), even some that unexpectedly did not match reported assignments^9,26^; namely, G288, F289, G400, and F401.

The ^1^H-^15^N HSQC spectrum of TDP-43 PrLD displayed poor chemical shift dispersion in the ^1^HN dimension, consistent with a disordered domain (**Fig. 1a**), and the NMR assignments confirmed the presence of the α-helix in the central region of TDP-43 PrLD (at residues 320-340) (**Fig. 1b**). In addition to the helical region, Li *et al*. have shown that aromatic residues within the GAroS segments outside the central region are also relevant to some extent to the assembly process^11^. In order to interrogate the role of the helical component and the GAroS regions in the intrinsic aggregation of TDP-43 PrLD at pH 4, we analyzed peak intensities of the ^1^H-^15^N HSQC spectrum. This strategy has been used to characterize the helix-driven aggregation pathway of TDP-43 PrLD upon droplet induction with salt at higher pH values^10,11^. The peak intensities observed for the first ^1^H-^15^N HSQC spectra recorded within the first hour from sample preparation at two different concentrations (55 and 110 µM) revealed non-homogeneous peak intensities throughout the PrLD sequence, with reduced intensity for the central residues containing the helical segment (**Fig. 1b**, black bars), which is interpreted as shift from the monomeric towards the aggregate state mediated by helix-helix intermolecular interactions, in line with previous observations^9-12^. Over the course of *ca*. 20 hours, the greater signal intensity loss of the central region became more evident at both concentrations (**Fig. 1b**, red bars). A closer inspection to these data revealed that, in addition to the central region containing the helix, other signals show reductions in signal intensity. Although these are minor changes, they are consistently observed over time and at the two concentrations (55 and 110 µM) as lower-than-average intensity values, and mostly map to the two GAroS segments that are N- (residues 267-310) and C-terminal (361-414) to the central 311-360 region (**Fig. 1b**, dashed horizontal lines). In particular, we highlight the following stretches of hydrophobic residues: four Phe-Gly-rich motifs at positions 276-284, 288-290, 367-368, and 396-403; one Tyr-Ser-Gly motif at 373-376; and two Trp-Gly motifs at positions 384-386 and 411-413 (**Fig. 1b, 1c**). This result correlates very well with mutational analyses that showed that, in addition to the helical region, Phe283, Phe289 (in the GAroS segment that is N-terminal to the helix) and Tyr374, Trp385, Phe401, and Trp412 (in the GAroS C-terminal to the helix) are required for droplet induction, since their mutation to glycines abolish salt-induced aggregation^11^. On this basis, we reasoned that the intrinsic aggregation we observed here does not follow an alternative mechanism with respect to that firmly established in the literature^9-12^.

**Fig. 1.**
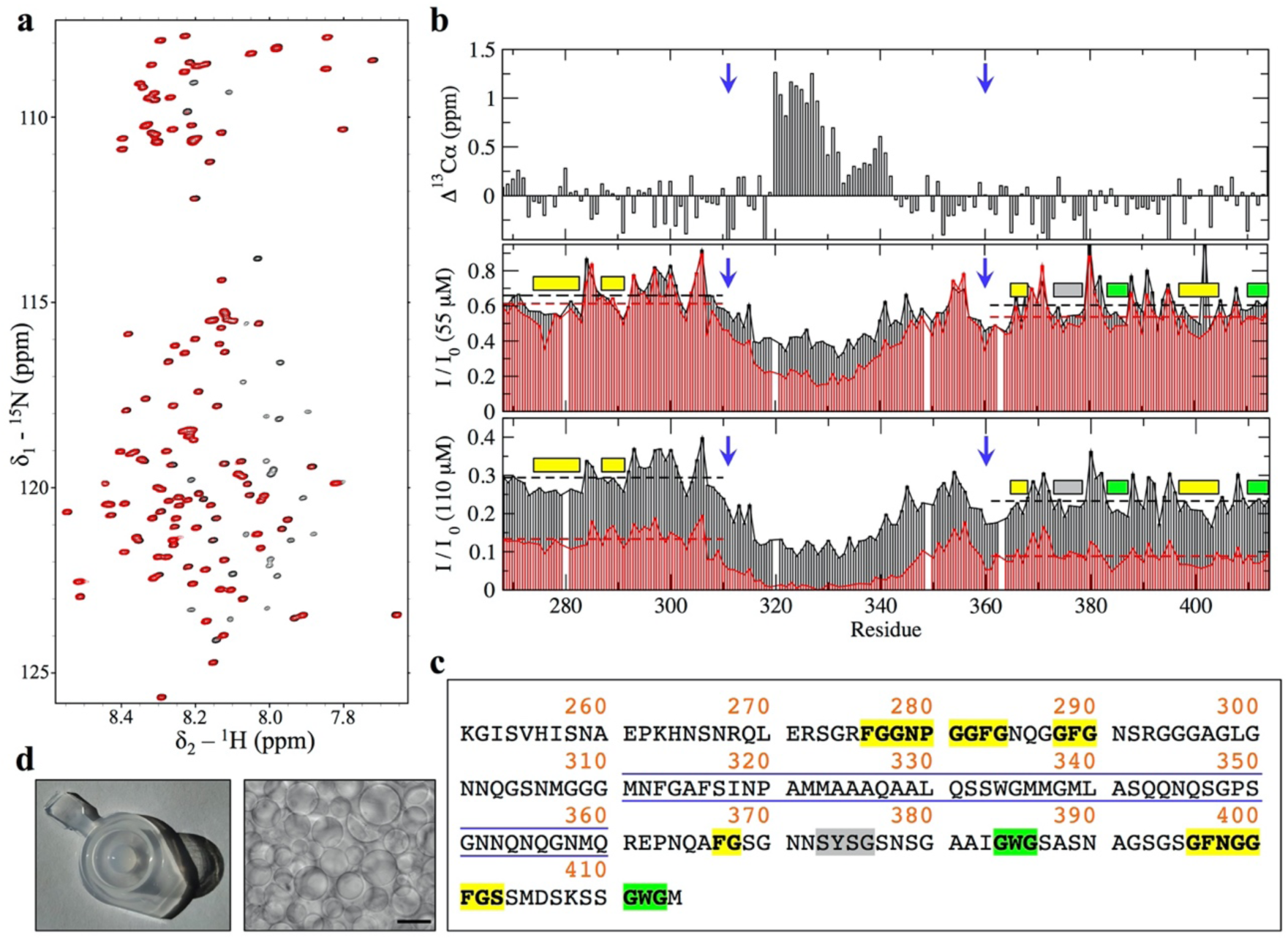
LSNMR characterization of the PrLD of TDP-43’s aggregation at low pH. **a** The ^1^H-^15^N HSQC spectra of TDP-43 PrLD at pH 4 and 25°C (110 µM), showing a narrow ^1^H amide chemical shift range typical of a disordered region. Black peaks correspond to a spectrum recorded within one hour from sample preparation, the time elapsed between elution from the column and the beginning of the NMR experiments (see Methods). The spectrum in red was recorded at the end of the NMR experiments recorded on this sample (**Table S2**), where signals corresponding to some but not all residues disappear due to self-association. **b** Signals whose intensity is lost over time correspond to the central region spanning residues 311-360 (delimited by blue arrows), which contain an α-helix as evinced by ^13^Cα chemical shifts (top panel). In addition, several short segments within the two regions N- and C-terminal to the 311-360 core (*i.e*., 267-310 and 361-414) also exhibit moderate signal loss, which are consistently observed at 55 µM (middle panel) and at 110 µM (bottom panel). This is illustrated by plotting the intensity values from the first ^1^H-^15^N HSQC (I_0_, black bars, one hour from sample preparation), and those from a ^1^H-^15^N HSQC recorded ∼20 h later, plotted as I/I_0_ (red bars, I and I_0_ correspond to intensity values for the second and first ^1^H-^15^N HSQC, respectively). Note that average values for the two regions 267-310 and 361-414 N- and C-terminal to the central segment are shown as dashed horizontal lines (black and red colors for the I_0_ and I/I_0_ data sets, respectively), which highlights that the regions with lower-than-the-average intensities mostly map to segments containing aromatic residues: Phe (yellow boxes), Tyr (gray box) and Trp (green boxes). **c** Amino acid sequence of TDP-43’s PrLD illustrating the central region composed of residues 311-360 (blue lines, to match blue arrows in panel b), which was the focus of some earlier studies and contains the helical segment important for assembly, as well as the aromatic residues outside this core (Phe in yellow, Tyr in gray, Trp in green) **d** At the end of the NMR experiments, the samples were found aggregated (left panel), and displayed extensive droplet formation (right panel, bar scale 100 μm). This was imaged on the 110 µM sample.

After ∼7 (at 55 µM) or ∼3.5 days (at 110 µM), the signals corresponding to residues within the central region and GAroS segments disappeared (**Fig. 1a**, red spectrum), and we observed that the NMR samples contained aggregated material displaying extensive droplet formation (**Fig. 1d**). Overall, these findings evince that: (i) the TDP-43 PrLD self-assembles at low pH and without salt, conditions where it was reported to not aggregate^13,14^, and that (ii) the aggregation process follows a helix-driven, GAroS-assisted mechanism of phase separation into droplets similar to that reported under droplet induction by salt or RNAs^9-12^. We interpret these observations as an *intrinsic* aggregation pathway of TDP-43 PrLD that is solely encoded by its amino acid sequence. Thus, it may well manifest over a broad range of conditions in cells, which would lead to the formation of relevant aggregate species at pH values below those normally considered physiological but relevant for metabolic stress conditions^5^ and lysosome interiors^27^. Considering that all the prior work that uncovered the mechanism of droplet assembly did not detect fibrils (*i.e*., droplets were endpoint species following TDP-43 PrLD aggregation)^9-12^, we next interrogated whether the droplets we detect could evolve into amyloid-like assemblies.

### The TDP-43 PrLD droplet environment outbursts amyloid fibrils at the droplet surface/solvent interface

Following the discussion of droplet versus fibrils as end point of distinct aggregation pathways, it has been very recently proposed that droplet assembly in FUS, hnRNPA1, and potentially most archetypical PrLDs, is encoded by its primary sequence as enabled by a symmetric distribution of aromatic residues. More precisely, this refers to aromatic Phe, Trp, or Tyr “stickers” residues separated from each other by stretches of 3 to 12 “spacers” residues, such as Ser or Gly^28^. Within this framework, such symmetric distribution is not fulfilled if the aromatic stickers cluster together in the primary sequence (*i.e*., without interspaced residues), and this causes irreversible fibrillization instead over droplet formation^28^. We identified such a “sticker-and-spacers” symmetric distribution in the GAroS regions of TDP-43 PrLD, which may explain the intrinsic aggregation that we proposed to be encoded solely by its primary sequence. More precisely, the N-terminal region to the α-helix contains three Phe “sticker” residues, Phe276, Phe283, and Phe289, which are separated by two stretches of “spacers” spanning six (F_276_GGPNGGF_283_) and five (F_283_GNQGGF_289_) residues, respectively (**Fig. 1c**). Similarly, the GAroS region that is C-terminal to the α-helix contains six aromatic residues, which can be grouped in the following sticker-and-spacers pairs: Tyr374 - Trp385 (separated by 10 spacers: Y_374_SGSNSGAAIGW_385_), Trp385 - Phe397 (separated by 11 spacers: W_385_GSASNAGSGSGF_397_), Phe397 - Phe401 (separated by 3 spacers: F_397_NGGF_401_), and Phe401 - Trp412 (separated by 10 spacers: F_401_GSSMDSKSSGW_412_) (**Fig. 1c**). Therefore, our interpretation that the lower peak intensities in the ^1^H-^15^N HSQC mapping to the GAroS regions resulted from intermolecular interactions driving the monomer to droplet transition (**Fig. 1b**) fits well within this *stickers-and-spacers* model for PrLD-containing proteins.

One distinctive feature between TDP-43 PrLD and other PrLDs is the helical region; its tendency to self-aggregate not only reduces the number of “stickers” required for droplet formation^11^, but also drives the assembly of amyloid fibrils in distinct aggregation pathways^13,14^. On this basis, and to test the model of uniform distribution of aromatic stickers as a driver for LLPS, but not amyloid formation^28^, we explored whether the droplets formed after the LSNMR experiments (**Fig. 1d**) lead to the formation of amyloid fibrils. To this end, we monitored Thioflavin T (ThT) fluorescence using confocal microscopy. We observed a strong enhancement of ThT fluorescence in the droplets formed in the two samples at 55 and 110 µM (**Fig. 2a-2c**) and, consistent with this result, dense networks of fibrils were imaged by transmission electron microscopy (TEM) (**Fig. 2d**). The observation of ThT-reactive droplets in two distinct samples that aggregated following the same mechanism, as uncovered by LSNMR (**Fig. 1b**), together the imaging of the fibrils by TEM, firmly establishes that TDP-43 PrLD fibril formation is coupled to droplets under these conditions.

**Fig. 2.**
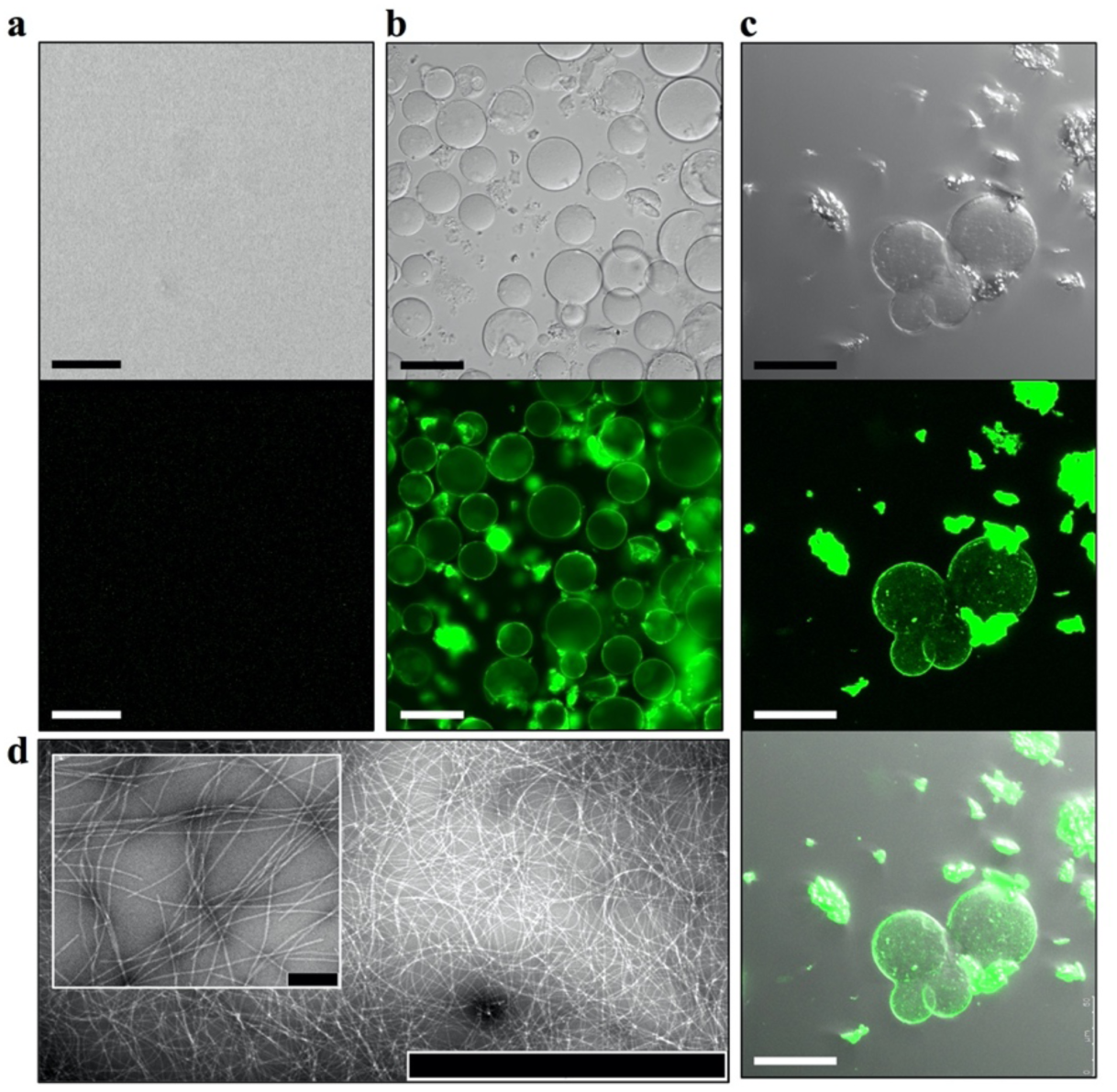
Droplet assembly is coupled to fibril formation. **a** The TDP-43’s PrLD sample eluted from the column is free of droplets (top). To test for the presence of amyloid fibrils, Thioflavin T (ThT) was added to the sample and visualized by fluorescence microscopy (bottom). The lack of fluorescence indicates that no amyloid fibrils are present at the beginning of the experiments. **b** Aggregated samples (*i.e*., samples where no signal is detected in LSNMR experiments) present extensive droplet formation, as well as irregularly shaped clumps (top, light transmitted image). Interestingly, both the droplets and the clumps were recognized by ThT (bottom, fluorescent image). These images were collected using fluorescence microscopy on 110 µM TDP-43 PrLD. The scale bars in *a* and *b* are 100 µm. **c** Confocal microscopy images of the 55 µM aggregated sample (transmitted light, top; fluorescence, middle; overlay, bottom) showing ThT bound to droplets surface as well as to the irregularly shaped clamps, suggesting that the latter are made by amyloid fibrils which are formed and released from the droplet surface. Scale bar is 50 µm. **d** TEM micrographs confirmed the presence of amyloid fibrils, in agreement with ThT fluorescence (scale bar: 2 µm; at inset image: 200 nm).

The above results evoke the recent work by Gui *et al*., who used ThT fluorescence and confocal microscopy to provide direct visual evidence that fibrils from hnRNPA1 –another protein with a PrLD– formed from induced protein droplets^16^. A close inspection to both hnRNPA1 in Gui *et al*.’s paper, and TDP-43 PrLD in the present work, revealed a critical difference: whereas ThT fluorescence shines from the droplet interior in hnRNPA1 aggregates, indicative of fibrils confined within the droplet interior, the TDP-43 PrLD fibrils seems to be present at the surface, but not inside the droplet. One distinctive advantage of confocal microscopy is the possibility of scanning the entire volume of a droplet, slice by slice, and this feature allowed us to look in closer detail at TDP-43 PrLD droplets. In **Fig. 3**, we provide direct visualization that fibrils are in the surface and not in the interior of the droplet. Furthermore, they seem to detach upon reaching a critical size (**Fig. 3**). Based on these differences between hnRNPA1 versus TDP-43 PrLD (fibrillization in the droplet interior versus the droplet surface, respectively), direct visualization of ThT fluorescence of proteins in condensates represents a promising approach to scrutinize details of fibril formation in droplet environments.

**Fig 3.**
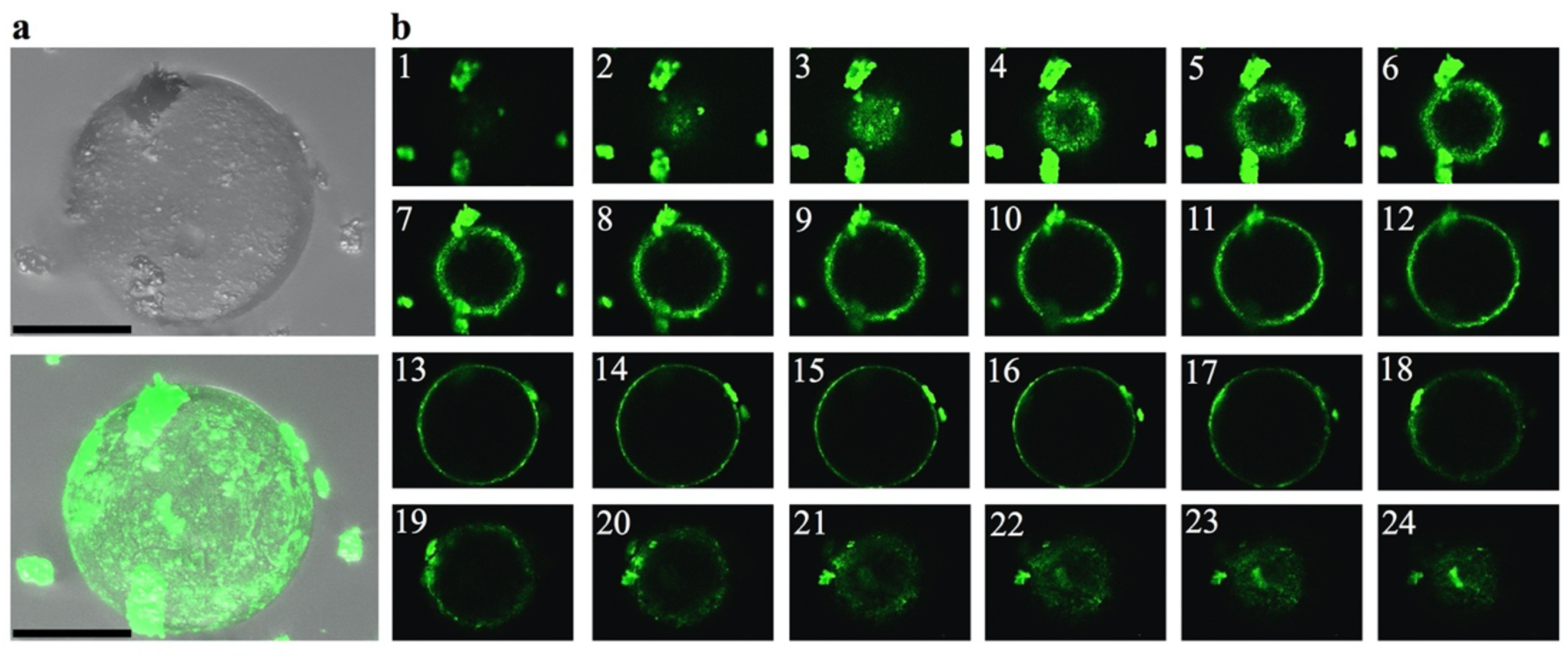
ThT-positive, TDP-43’s PrLD fibrils emerge from protein droplets. **a** Visualization of a ThT-stained droplet from the 110 µM aggregated sample by confocal microscopy (top: transmitted light image, bottom: overlay with the ThT fluorescent image. Scale bar: 50 µM). The non-homogeneous staining suggests that fibril assembly occurs on droplet surface, and not its interior. Scale bar is 50 µm. **b** Sections of the droplet evince that the droplet interior is devoid of fibrils, whereas ThT fluorescence maps the droplet surface, at the droplet/solvent interface boundaries, and suggests detachment of fibrils once a certain size is reached. Images 1 to 24 are selected sections from a collection of 42 slices passing from pole to equator to pole that illustrate the distribution of ThT-positive aggregates across the droplet surface.

### The SSNMR signature of TDP-43 PrLD fibrils reveals an amyloid core that builds on aromatic stickers

We have shown that, at low pH, TDP-43 PrLD has an intrinsic ability to assemble droplets by virtue of its central region and the GAroS segments with aromatic strickers and spacers (**Fig. 1**). We have then shown that this intrinsic aggregation pathway couples droplet formation with the accumulation of amyloid fibrils at the droplet surface/solvent interface. What structural factors drive the conversion of fibrils? During droplet assembly, Trp-Gly motifs are the most important residues in assisting the helical region^9,11^. In fibrils, previous cryo-EM and SSNMR studies with the TDP-43 PrLD (311-360) peptides proved that the helical region suffices to form amyloid fibrils^15,19^. However, these peptides lack the GAroS regions (residues 267-310 and 361-414) and do not inform on their participation in the fibrillar structures, as they do in droplets^9,11^ (**Fig. 1**). Thus, we ought to explore whether the symmetrically distributed aromatic residues within the TDP-43 PrLD GAroS would form part of the fibril core, by taking advantage of their unique structural signature in the NMR spectra.

We analyzed the aggregated material that became invisible to LSNMR by means of SSNMR. Although full structure determination by SSNMR requires multiple samples with different labeling schemes and in large quantities^29^, we could reconstitute a modest amount of the fibril conformation that formed at the droplet/solvent interface. Cross-polarization Magic Angle Spinning- (CPMAS)-based experiments are sensitive to molecular mobility, allowing us to scrutinize the presence of aromatic residues immobilized within the fibril core. **Figure 4a** shows a ^13^C-^13^C spectrum recorded under appropriate conditions to ensure detection of sequential residues (*i.e*., j and j+1) that remain static and rigid. No Trp residues could be detected in the characteristic region corresponding to aromatic ^13^C nuclei (*i.e*. no indole side chains are observed). This result is curious because Trp residues are key to droplet formation by the TDP-43 PrLD^9,11^, such that their mutation to Gly abolish droplet initiation^11^. One potential explanation could be that Trp residues are exposed in the fibril, and not fixed in the core interior, such that their indole rings are on average less rigid and do not contribute strong signals in CPMAS-based experiments. However, the most striking observation is that we identified sequential contacts that correspond to several Phe/Tyr, Gly and residues (**Fig. 4a**), which unambiguously indicate that these motifs are immobilized within a fibrillar core containing β-rich segments, as evinced by distinctive cross-peaks in the Cα/Cβ serine region (**Fig. 4a**). This is reminiscent of structural models of other PrLD-containing fibrils, in which the amyloid core is formed by Ser/Gly/Aromatic-rich motifs^30^. Unlike Trp residues, mutation of Phe residues to Gly did not abolish droplet initiation, as long as the Trp residues remained unchanged^11^. This observation indicates that Phe-Gly motifs have distinct roles in the droplet and the fibril forms. While a structure determination is still in progress, for which we are optimizing the production of larger amount of samples with distinct labeling schemes, the 1D slices of the 2D dataset already indicate that the signal from Phe/Tyr aromatic ^13^C, Gly’s ^13^Cα, and Ser’s ^13^Cβ nuclei are unambiguously detectable under our experimental conditions (**Fig. 4b**). When the same experiment is recorded with a shorter mixing time, only intra-residual correlations are observed, supporting the build-up of sequential connectivities as the mixing time is increased (**Fig. 4c**). The GAroS regions of TDP-43 PrLD contains six Phe-Gly motifs, three of which exist as Phe-Gly-Ser triads (Phe367, Phe397, and Phe401) (**Fig. 1c)**. There is also just one Tyr residue (Tyr374), which appears as Tyr-Ser-Gly (**Fig. 1c)**. Tentatively, it seems reasonable to speculate that these motifs may be part of the structural core.

**Fig. 4.**
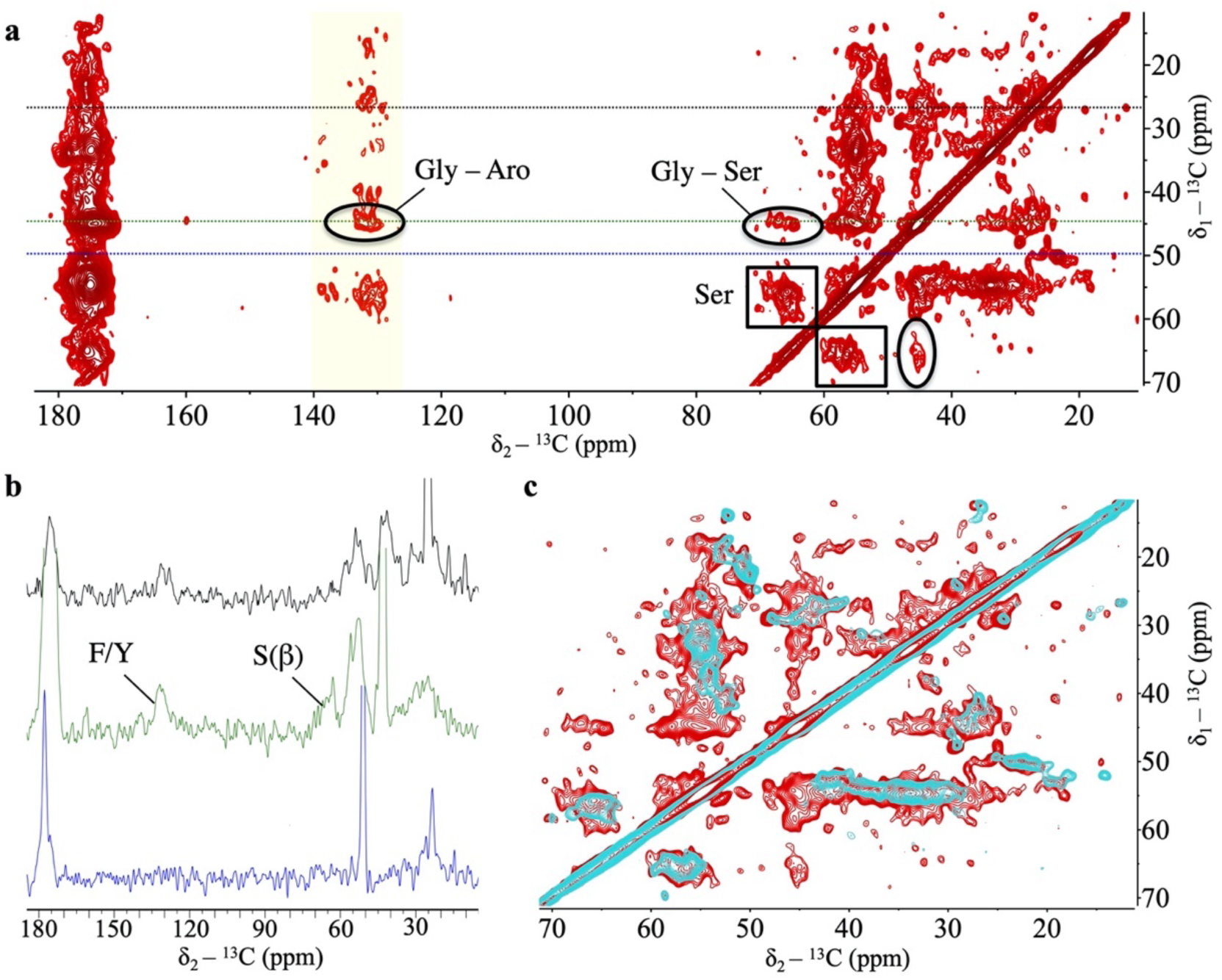
Fibrils from the PrLD of TDP-43 are stabilized by Phe-Gly motifs. **a** 2D ^13^C-^13^C DARR spectrum recorded with a 250 ms mixing time on fibrils formed by the PrLD of TDP-43. This experiment detects residues immobilized within the fibril core. Fibrils are rich in β-sheet content, as indicated by the presence of Ser residues showing differences of ^13^Cβ -^13^Cα > 8 ppm (black boxes). Cross-peaks between Ser and Gly (black ellipses) indicate that Ser-Gly motifs, which are located outside the helical region, are relevant in the fibril form. Signals around 130-140 ppm (yellow shaded region) correspond to aromatic ^13^C from Phe/Tyr residues. Considering that there is only one Tyr in the PrLD sequence, most cross-peaks arise from Phe (^13^Cδ/^13^Cε) – Gly (^13^Cα) (black circles). **b**, Horizontal slices extracted from the 250ms DARR shown in *a*, taken at: δ_1_= 27.03 ppm (in black), which illustrates a crowded region that covers aliphatic, aromatic and carbonyl peaks); δ_1_= 45.07 ppm (in green), which is distinctive of Gly ^13^Cα chemical shift values, illustrating signals contributed by aromatic ^13^C nuclei from Phe/Tyr, as well as by ^13^Cβ of Ser, as identified by their distinctive chemical shift values. This supports the presence of immobilized Gly, Ser, and Phe residues in the fibril core; and δ_1_= 49.69 ppm (in blue), which lacks strong resonances thus facilitating an appreciation of the overall signal-to-noise of the SSNMR data. **c** A short mixing time (20 ms) 2D ^13^C-^13^C DARR spectrum (in cyan) overlapped onto the 250 ms spectrum from panel *a* (in red). At this short mixing, only intra-residue contacts are detected, which supports the assignment of sequential contacts that build up at larger mixing times.

## Discussion

To date, several groups have presented studies of intrinsically disordered protein domains undergoing phase separation, wherein the microdroplets formed eventually evolve into amyloids^14-18^. Until now, it has been generally assumed that the residues responsible for forming the microdroplet are also those that become amyloid^13,15,31^. Here, we have combined LSNMR and SSNMR to obtain atomic level information on both processes. In contrast to previous expectations, we have uncovered that certain residues are responsible for droplet formation whereas others are key for amyloidogenesis, including aromatic “sticker” residues proposed to enable reversible droplet formation and not fibrillization^28^. This discovery has important implications for our fundamental understanding of these processes. Excitingly, it means that we should be able to develop inhibitors or modulators that target the residues that are key for one process (*i.e*. the Phe residues which promote amyloidogenesis), without affecting the other residues which promote physiological droplet formation to confront the stress situation. This is also supported by the observation that TDP-43 PrLD also assembles droplets that do not evolve into fibrils^9-12^. Importantly, these features could not be recruited in model fibrils from the 311-360 segments that exclude the GAroS segments harboring the Phe-Gly motifs (residues 267-310 and 361-414).

We postulate that the fibril conformation that forms at pH 4 may represent a pro-pathological structure resulting from the persistence of a droplet state, and that it may exist over a broad range of conditions based on the low content of titratable residues in the TPD-43 PrLD. This hypothesis is sustained by an intrinsic mechanism of droplet assembly, which remains essentially identical at pH 7 and in the presence of salts or RNA, and at pH4 without salts or RNA. Recently, Shenoy *et al*. carried out SSNMR measurements on TDP-43 PrLD fibrils assembled at pH 7.5^32^ and, interestingly, they did not report the Ser, Gly, and Phe/Tyr signals that we detect under similar mixing time conditions in our 2D ^13^C-^13^C spectra from fibrils assembled at pH 4 (**Fig. 2**). This suggests the reconstitution of alternative structures in the two studies, which we explain based on the droplet stage reported, rather than different pH conditions. The droplet environment enabled fibrillization at the droplet surface/solvent interface, and such fibril form was amplified by seeding soluble protein, as this strategy is commonly used to propagate selected amyloid conformations^33-35^. Shenoy *et al*. used a 4-mm rotor in their SSNMR study, for which they successfully prepared a large amount of fibrillar material, but did not inform the formation of droplets^32^. This is reminiscent of the different outcomes observed in cryo-EM and SSNMR studies on fibrils from the TDP-43 PrLD (311-360) segment, in which the fibrils that form from droplets revealed spectral signatures not compatible with the cryo-EM structures^15,19^. These observations in aggregate supports the view that TDP-43 PrLD participates in distinct aggregation pathways, with different end point species that include: (i) droplets without fibrils^9-12,19^, (ii) fibrils without droplets^13,32^, and (iii) droplets and fibrils^14,15^, as interpreted with the data informed to date. The droplet to fibril transitions are of potential relevance, as cytoplasmic TDP-43 droplets were shown to sequestrate components of the nuclear pore^36^, and co-aggregate formation between TDP-43 and nuclear pore components emerged as a hallmark of ALS/FTD^37^, and they may well be hybrid amyloids similar to the RIPK1-RIPK3 necrosome core^29^. Amidst these diverse findings, ours are the first to directly link droplets to amyloid formation by combining LSNMR and SSNMR. Deciphering the structural features of the various TDP-43 aggregate species in diverse contexts^3,21^, such as the Phe-Gly motifs shown here to create a fibril core is an important step towards understanding physiological versus pathological aggregation.

## Methods

### Protein expression and purification

The N-terminally hexa-His-tagged PrLD construct corresponding to TDP-43(267-414) was a gift from Prof. Nicolas Fawzi (Addgene plasmid # 98669). Proteins were overexpressed in *E. coli* BL21 Star (DE3) (Life Technologies). Uniform ^15^N and ^15^N/^13^C labeling of TDP-43(267-414) was achieved by overexpression in M9 minimal media supplemented with ^15^N ammonium chloride and ^13^C glucose as the sole sources of nitrogen and carbon, respectively. *E. coli* BL21 Star (DE3) carrying TDP-43(267-414) were pre-inoculated overnight at 37°C. The day after, 200 ml of M9 minimal media were added to 4 mL of pre-inoculum and the cell growth was monitored using Biophotometer D30 (Eppendorf, Hamburg, Germany) until the growth rate was 0.8 (OD at 600 nm). Then, the protein expression was induced overnight at 30°C by adding 1 mM IPTG.

Cells were collected by centrifugation at 7,000 × g for 1 hour 4°C and the resulting pellet was resuspended in 30 mL lysis buffer (50 mM NaH_2_PO_4_, 300 mM NaCl, 10 mM Imidazole pH 8.0) supplemented with one tablet EDTA-free protease inhibitor cocktail (Roche Diagnostics) and 1 mM PMSF (SigmaAldrich, St Louis, MO, USA). Cell lysis were performed using EmulsiFlex-C3 (Avestin Europe GmbH, Mannheim, Germany) and Bioruptor UCD-200 (Diagenode, Belgium). Inclusion bodies were recovered by centrifugation of cell lysate at 7,000 x g for 1 hour at 4°C and were resuspended in 20 mL denaturing binding buffer (20 mM Tris-Cl, 500 mM NaCl, 10 mM Imidazole, 1 mM DTT, 8 M urea, pH 8.0). Further centrifugation at 7,000 × g for 1 hour, 4°C was performed in order to pellet any remaining insoluble cell debris.

Ni-NTA Agarose beads (Qiagen Inc, Gaithersburg, MD, USA) were used to bind the target protein, and then were washed three times in 15 mL of denaturing binding buffer and each washing step was interspersed with centrifugation at 500 × g for 5 minutes, RT. Beads were then incubated with cell lysate for 4 hours rotating and successively collected by centrifugation at 500 × g for 5 minutes, RT. Finally, beads were washed three times in 2 ml of denaturing binding buffer and protein was eluted with imidazole gradient buffers, as described below:

1. 1 ml 20 mM Tris-Cl, 500 mM NaCl, 250 mM imidazole, 1 mM DTT, 8 M urea, with rotation of 2 hours at RT.
2. 1ml, 20 mM Tris-Cl, 500 mM NaCl, 300mM imidazole, 1 mM DTT, 8 M urea, with rotation overnight at 4°C.
3. 1ml 20 mM Tris-Cl, 500 mM NaCl, 300mM imidazole, 1 mM DTT, 8 M urea, with rotation of 2 hours at RT.
4. 1ml 20 mM Tris-Cl, 500 mM NaCl, 500 mM imidazole, 1 mM DTT, 8 M urea, with rotation of 6 hours at RT.

Each elution step was interspersed with centrifugation at 500 ×g for 5 minutes, RT, and from the total elution pool ∼3 mg of protein are obtained.

The pH of the samples from the TDP-43 PrLD constructs was then lowered to 4.0, and applied to a PD-10 gel filtration column (GE Healthcare Ltd, Little Chalfont, UK), which had been previously pre-equilibrated with the LSNMR buffer: 1 mM CD_3_COOD in 85/15 H_2_O/D_2_O. The eluted fractions containing the PrLD construct, as identified by UV absorbance, were then concentrated to ∼550 µL using an ultrafiltration device (Amicon Corp., Danvers, MA, USA). The final urea concentration was less than 12 mM as determined by refractive index.

### Choice of the best LSNMR assignment strategy

The TDP-43 PrLD is intrinsically disordered and low-complexity (*i.e*., high redundancy in the amino acid sequence), and therefore poorly dispersed amide ^1^H peaks and clustering of the ^13^Cα and ^13^Cβ chemical shifts around the random coil values are anticipated to complicate the assignment process. On the other hand, methods based on ^13^C detection exploiting that ^15^N and ^13^CO signals remain well dispersed in disordered proteins cannot be applied to the TDP-43 PrLD, as these methods require high protein concentration conditions to overcome the low ^13^CO sensitivity, under which TDP-43 PrLD quickly aggregates. With these limitations ahead, we resorted to ^1^H-detected methods that mainly consist in connecting two consecutive NH groups through their correlations with one or more of the ^13^C spins located between them: ^13^Cα, ^13^Cβ and ^13^CO. Due to the limitations mentioned earlier, we reasoned that six 3D experiments would be required to unambiguously obtain backbone chemical shifts; namely, (i) HNCO, (ii) HN(CA)CO, (iii) HNCA, (iv) HN(CO)CA, (v) CBCA(CO)NH, and (vi) HNCACB. Recording this set of experiments with an acceptable resolution entails an average time of ∼90 hours, as illustrated in **Table S1**, which is by far too long for the TDP-43 PrLD to remain in solution due to its high aggregation tendency.

We then reasoned what would be the best strategy to accomplish the NMR assignments of our protein, ruling out the use of non-uniform sampling (NUS) schemes as the low concentration conditions the percentage of sampling that would be reduced may not represent a great time gain. With all these drawbacks, we proposed using a different strategy in which just three 3D experiments enabled unambiguous assignments of all signals in the ^1^H-^15^N HSQC spectrum of TDP-43 PrLD (**Table S2**). The approach consists of two experiments, one H(NCOCA)HN and one (H)N(COCA)NH, that allowed us to directly connect consecutive amide groups from all signals in the ^1^H-^15^N HSQC. Using the information from a CBCA(CO)NH experiment, the spin systems are well defined through the ^13^Cα and ^13^Cβ chemical shifts. With this strategy, experiments are acquired in a time of ∼55 hours, that is to say, 37.5% less than the conventional triple resonance strategy described above. These experiments were recorded on the 110 µM sample, along with an additional HNCA to corroborate the assignments and to obtain the ^13^CA chemical shifts for the four residues preceding prolines (P280, P320, P349, and P363), and for the last residue M414. The experiment time of the HNCA was 5.5 hours (**Table S2**), which still represent a reduction of 31% with respect to the conventional approach listed in Table S1. Finally, a CC(CO)NH experiment was recorded on this sample (experiment time 22.5 hours) to assign side chain ^13^C nuclei from Gln, Arg, Lys, Pro, Ile, and Leu residues. After these ∼3.5 days of NMR measurements, some signals in the ^1^H-^15^N HSQC are missing and the sample was aggregated. To report the full set of backbone assignments, the ^13^CO and ^1^Hα chemical shifts were obtained by recording a HNCO (5.5 hours) and a HBHA(CO)NH (33,2 hours) on the 55 µM sample. The latter experiment also provides the assignments for ^1^Hβ nuclei.

### LSNMR experiments

The above liquid state NMR experiments were collected at pH 4.0 and 298 K on a Bruker AV-800 US2 (800MHz ^1^H frequency) spectrometer equipped with a cryoprobe and Z-gradients. The two samples (55 and 110 µM) were prepared by concentration of the eluted fractions to ∼550 µL, from which 300 µL of were transferred into D_2_O-matched, 5 mm Shigemi tubes. To obtain the ^13^CO, ^13^Cα, ^13^Cβ, ^15^N, ^1^HN, ^1^Hα and ^1^Hβ NMR assignments following the strategy presented in the previous section, we started from a root ^1^H-^15^N HSQC spectrum, which served as the first spectrum to carry out the peak intensity analyses in Fig. 1. This experiment was measured again after ∼20 hours with the following parameters: 4 scans, 12 and 20 ppm as spectral widths for ^1^H and ^15^N, respectively, and transmitter frequency offsets of 4.75 and 116.5 ppm for ^1^H and ^15^N, respectively. The remaining experiments to obtain chemical shifts to be deposited in the Biological Magnetic Resonance Bank were recorded with the following parameters (see also **Table S1** and **S2**):

CBCA(CO)NH (8 scans, 12, 20, and 75 ppm as ^1^H, ^15^N, and ^13^C spectral widths, respectively, and transmitter frequency offsets of 4.75, 116.5, and 44 ppm for ^1^H, ^15^N, and ^13^C, respectively); H(NCOCA)HN (12 scans, 12, 20, and 4.25 ppm as ^1^H, ^15^N, and ^1^H spectral widths, respectively, and transmitter frequency offsets of 4.75, 116.5, and 7.125 ppm for ^1^H, ^15^N, and ^1^H, respectively); (H)N(COCA)NH (12 scans, 12, 20, and 20 ppm as ^1^H, ^15^N, and ^1^H spectral widths, respectively, and transmitter frequency offsets of 4.75, 116.5, and 116.5 ppm for ^1^H, ^15^N, and ^15^N, respectively); HNCA (4 scans, 12, 20, and 30 ppm as ^1^H, ^15^N, and ^13^C spectral widths, respectively, and transmitter frequency offsets of 4.75, 116.5, and 54 ppm for ^1^H, ^15^N, and ^13^C, respectively); HNCO (8 scans, 12, 20, and 12 ppm as ^1^H, ^15^N, and ^13^C spectral widths, respectively, and transmitter frequency offsets of 4.75, 116.5, and 174 ppm for ^1^H, ^15^N, and ^13^C, respectively); HBHA(CO)NH (12 scans, 12, 20, and 6, ppm as ^1^H, ^15^N, and ^1^H spectral widths, respectively, and transmitter frequency offsets of 4.75, 116.5, and 4.75 ppm for ^1^H, ^15^N, and ^1^H, respectively), and a CC(CO)NH experiment (8 scans, 12, 20, and 75 ppm as ^1^H, ^15^N, and ^13^C spectral widths, respectively, and transmitter frequency offsets of 4.75, 116.5, and 39 ppm for ^1^H, ^15^N, and ^13^C, respectively)

Proton chemical shifts were directly referenced using DSS on a TDP-43 PrLD sample prepared for this purpose, and ^13^C and ^15^N chemical shifts chemical shifts were referenced indirectly. Conformational chemical shifts, Δ(^13^Cα), where calculated as d^13^Cα(exp) -δ^13^Cα(ref), with δ^13^Cα(exp) being our measured chemical shifts and δ^13^Cα(ref) reference values obtained using the sequence of the TDP-43 PrLD construct and the data compiled by Poulsen, Dyson, and co-workers, at 25 °C and pH 4, as implemented in the chemical shift calculator http://spin.niddk.nih.gov/bax/nmrserver/Poulsen_rc_CS^39-41^.

All spectra were processed using either NMRPipe^42^ or Topspin 4.0.8 (Bruker Biospin, Germany), and peak-picking and spectral assignment was conducted using NMRFAM-Sparky^43^. The NMR chemical shifts are deposited in the Biological Magnetic Resonance Bank (BMRB) under accession code 50154.

### SSNMR experiments

Solid-state NMR experiments were recorded on the aggregated material that formed following droplet formation during our LSNMR studies, which was amplified by seeding soluble protein prepared as described above. As a reference, from the most concentrated LSNMR samples (110 µM), the 300 µL from the Shigemi tube contains approximately 600 µg of material, assuming that 100% of the protein converted into fibrils when no signal is detected. After three weeks of incubation, the supernatant was carefully separated from the aggregated material settled on the bottom of the Eppendorf, which was lyophilized and transferred into a rotor using Bruker MAS rotor tools. The experiments were conducted on a Bruker 17.6 T spectrometer (750 MHz ^1^H frequency) using an HCN 1.3 mm MAS probe. 2D ^13^C-^13^C Dipolar Assisted Rotational Resonance (DARR) spectrum^44^ were recorded with mixing times of 20 and 250 ms, spinning the rotor containing the sample at a MAS rate of 17 and 20 kHz, respectively. The longer, 250 ms mixing time seemed appropriate to observe cross-peaks corresponding to sequential (j to j+1) residues, to detect correlations between Gly and Phe, Trp or Tyr, based on the characteristic chemical shifts of the Gly ^13^Cα and ^13^Caro chemical shift values distinctive for each type of aromatic residue. The spectra were recorded with the acquisition parameters listed in **Table S3**, 64 scans, using a spectral width of 220.9 ppm in the direct and indirect dimensions, with acquisition times of 20 and 2.5 ms, respectively, and setting the transmitter frequency offset to 100 ppm. The DARR spectrum was processed using Topspin 4.0.8, with chemical shifts referenced to DSS.

### Visualization of amyloid fibrils and protein droplets

Liquid droplets were directly visualized by spotting aliquots of the two (55 and 110 µM) samples onto glass coverslips using a Leica TCS SP2 inverted confocal microscope equipped with 7 laser lines, by both transmitted light (**Fig. 1d**) and fluorescence imaging (**Fig. 2-3**). The latter were obtained by addition of 1 μL of 1mM ThT and laser excitation at 457 nm, following the protocol by Gui *et al*. ^16^. To confirm that the ThT reactive species were amyloids, the samples were directly adsorbed onto carbon-coated 300-mesh copper grids, and negatively stained by incubation with 2% uranyl-acetate for direct visualization by transmission electron microscopy on a JEOL JEM-1011 electron microscope equipped with a TVIPS TemCam CMOS. Images acquired at a magnification of 30,000x and an accelerating voltage of 1,000 kV.

## Acknowledgments

This work has been funded by Grants CTQ2017-84371-P to D.P.-U. and SAF2016-76678-C2-2-R to D.V.L., from the Spanish MINECO; Grant MCB1412253 from the U. S. National Science Foundation to A.E.M; AriSLA (PathensTDP project) to E.B.; and Grant LCF/BQ/PR19/11700003 from La Caixa Foundation (ID 100010434) to M.M.

NMR experiments were performed in the “Manuel Rico” NMR Laboratory (LMR) of the Spanish National Research Council (CSIC), a node of the Spanish Large-Scale National Facility (ICTS R-LRB).

## References

1 Neumann, M. et al. Ubiquitinated TDP-43 in frontotemporal lobar degeneration and amyotrophic lateral sclerosis. Science 314, 130–133, doi:10.1126/science.1134108 (2006).

2 Nelson, P. T. et al. Limbic-predominant age-related TDP-43 encephalopathy (LATE): consensus working group report. Brain : a journal of neurology 142, 1503–1527, doi:10.1093/brain/awz099 (2019).

3 Vogler, T. O. et al. TDP-43 and RNA form amyloid-like myo-granules in regenerating muscle. Nature 563, 508–513, doi:10.1038/s41586-018-0665-2 (2018).

4 Chu, J. F., Majumder, P., Chatterjee, B., Huang, S. L. & Shen, C. J. TDP-43 Regulates Coupled Dendritic mRNA Transport-Translation Processes in Co-operation with FMRP and Staufen1. Cell reports 29, 3118–3133 e3116, doi:10.1016/j.celrep.2019.10.061 (2019).

5 Zhang, P. et al. Chronic optogenetic induction of stress granules is cytotoxic and reveals the evolution of ALS-FTD pathology. eLife 8, doi:10.7554/eLife.39578 (2019).

6 Mompean, M. et al. The TDP-43 N-terminal domain structure at high resolution. The FEBS journal 283, 1242–1260, doi:10.1111/febs.v13651 (2016).

7 Lukavsky, P. J. et al. Molecular basis of UG-rich RNA recognition by the human splicing factor TDP-43. Nature structural & molecular biology 20, 1443–1449, doi:10.1038/nsmb.2698 (2013).

8 Mompean, M., Baralle, M., Buratti, E. & Laurents, D. V. An Amyloid-Like Pathological Conformation of TDP-43 Is Stabilized by Hypercooperative Hydrogen Bonds. Frontiers in molecular neuroscience 9, 125, doi:10.3389/fnmol.2016.00125 (2016).

9 Conicella, A. E., Zerze, G. H., Mittal, J. & Fawzi, N. L. ALS Mutations Disrupt Phase Separation Mediated by alpha-Helical Structure in the TDP-43 Low-Complexity C-Terminal Domain. Structure 24, 1537–1549, doi:10.1016/j.str.2016.07.007 (2016).

10 Li, H. R. et al. The physical forces mediating self-association and phase-separation in the C-terminal domain of TDP-43. Biochimica et biophysica acta. Proteins and proteomics 1866, 214–223, doi:10.1016/j.bbapap.2017.10.001 (2018).

11 Li, H. R., Chiang, W. C., Chou, P. C., Wang, W. J. & Huang, J. R. TAR DNA-binding protein 43 (TDP-43) liquid-liquid phase separation is mediated by just a few aromatic residues. The Journal of biological chemistry 293, 6090–6098, doi:10.1074/jbc.AC117.001037 (2018).

12 Conicella, A. E. et al. TDP-43 alpha-helical structure tunes liquid-liquid phase separation and function. Proceedings of the National Academy of Sciences of the United States of America 117, 5883–5894, doi:10.1073/pnas.1912055117 (2020).

13 Lim, L., Wei, Y., Lu, Y. & Song, J. ALS-Causing Mutations Significantly Perturb the Self-Assembly and Interaction with Nucleic Acid of the Intrinsically Disordered Prion-Like Domain of TDP-43. PLoS biology 14, e1002338, doi:10.1371/journal.pbio.1002338 (2016).

14 Babinchak, W. M. et al. The role of liquid-liquid phase separation in aggregation of the TDP-43 low-complexity domain. The Journal of biological chemistry 294, 6306–6317, doi:10.1074/jbc.RA118.007222 (2019).

15 Zhuo, X. F. et al. Solid-State NMR Reveals the Structural Transformation of the TDP- 43 Amyloidogenic Region upon Fibrillation. Journal of the American Chemical Society, doi:10.1021/jacs.9b10736 (2020).

16 Gui, X. et al. Structural basis for reversible amyloids of hnRNPA1 elucidates their role in stress granule assembly. Nature communications 10, 2006, doi:10.1038/s41467-019-09902-7 (2019).

17 Molliex, A. et al. Phase separation by low complexity domains promotes stress granule assembly and drives pathological fibrillization. Cell 163, 123–133, doi:10.1016/j.cell.2015.09.015 (2015).

18 Patel, A. et al. A Liquid-to-Solid Phase Transition of the ALS Protein FUS Accelerated by Disease Mutation. Cell 162, 1066–1077, doi:10.1016/j.cell.2015.07.047 (2015).

19 Cao, Q., Boyer, D. R., Sawaya, M. R., Ge, P. & Eisenberg, D. S. Cryo-EM structures of four polymorphic TDP-43 amyloid cores. Nature structural & molecular biology 26, 619–627, doi:10.1038/s41594-019-0248-4 (2019).

20 Barbieri, E. M. & Shorter, J. TDP-43 shapeshifts to encipher FTD severity. Nature neuroscience 22, 3–5, doi:10.1038/s41593-018-0299-6 (2019).

21 Laferriere, F. et al. TDP-43 extracted from frontotemporal lobar degeneration subject brains displays distinct aggregate assemblies and neurotoxic effects reflecting disease progression rates. Nature neuroscience 22, 65–77, doi:10.1038/s41593-018-0294-y (2019).

22 Kroschwald, S. & Alberti, S. Gel or Die: Phase Separation as a Survival Strategy. Cell 168, 947–948, doi:10.1016/j.cell.2017.02.029 (2017).

23 Riback, J. A. et al. Stress-Triggered Phase Separation Is an Adaptive, Evolutionarily Tuned Response. Cell 168, 1028–1040 e1019, doi:10.1016/j.cell.2017.02.027 (2017).

24 van Leeuwen, W. & Rabouille, C. Cellular stress leads to the formation of membraneless stress assemblies in eukaryotic cells. Traffic 20, 623–638, doi:10.1111/tra.12669 (2019).

25 Asakawa, K., Handa, H. & Kawakami, K. Optogenetic modulation of TDP-43 oligomerization accelerates ALS-related pathologies in the spinal motor neurons. Nature communications 11, 1004, doi:10.1038/s41467-020-14815-x (2020).

26 Chen, T. S., Hsiao, C. L., Huang, S. J. & Huang, J. R. The nearest-neighbor effect on random-coil NMR chemical shifts demonstrated using a low-complexity amino-acid sequence. Protein Peptide Letters 23, 967–975, doi: 10.2174/0929866523666160920100045 (2016).

27 Settembre, C., Fraldi, A., Medina, D. L. & Ballabio, A. Signals from the lysosome: a control centre for cellular clearance and energy metabolism. Nature reviews. Molecular cell biology 14, 283–296, doi:10.1038/nrm3565 (2013).

28 Martin, E. W. et al. Valence and patterning of aromatic residues determine the phase behavior of prion-like domains. Science 367, 694–699, doi:10.1126/science.aaw8653 (2020).

29 Mompean, M. et al. The Structure of the Necrosome RIPK1-RIPK3 Core, a Human Hetero-Amyloid Signaling Complex. Cell 173, 1244–1253 e1210, doi:10.1016/j.cell.2018.03.032 (2018).

30 Murray, D. T. et al. Structure of FUS Protein Fibrils and Its Relevance to Self-Assembly and Phase Separation of Low-Complexity Domains. Cell 171, 615–627 e616, doi:10.1016/j.cell.2017.08.048 (2017).

31 Jiang, L. L. et al. Structural transformation of the amyloidogenic core region of TDP-43 protein initiates its aggregation and cytoplasmic inclusion. The Journal of biological chemistry 288, 19614–19624, doi:10.1074/jbc.M113.463828 (2013).

32 Shenoy, J. et al. Structural dissection of amyloid aggregates of TDP-43 and its C-terminal fragments TDP-35 and TDP-16. The FEBS journal, doi:10.1111/febs.15159 (2019).

33 Lu, J. X. et al. Molecular structure of beta-amyloid fibrils in Alzheimer’s disease brain tissue. Cell 154, 1257–1268, doi:10.1016/j.cell.2013.08.035 (2013).

34 Qiang, W., Yau, W. M., Lu, J. X., Collinge, J. & Tycko, R. Structural variation in amyloid-beta fibrils from Alzheimer’s disease clinical subtypes. Nature 541, 217–221, doi:10.1038/nature20814 (2017).

35 Walti, M. A. et al. Atomic-resolution structure of a disease-relevant Abeta(1-42) amyloid fibril. Proceedings of the National Academy of Sciences of the United States of America 113, E4976–4984, doi:10.1073/pnas.1600749113 (2016).

36 Gasset-Rosa, F. et al. Cytoplasmic TDP-43 De-mixing Independent of Stress Granules Drives Inhibition of Nuclear Import, Loss of Nuclear TDP-43, and Cell Death. Neuron 102, 339–357 e337, doi:10.1016/j.neuron.2019.02.038 (2019).

37 Chou, C. C. et al. TDP-43 pathology disrupts nuclear pore complexes and nucleocytoplasmic transport in ALS/FTD. Nature neuroscience 21, 228–239, doi:10.1038/s41593-017-0047-3 (2018).

38 Sonderby, I. E. et al. Dose response of the 16p11.2 distal copy number variant on intracranial volume and basal ganglia. Molecular psychiatry, doi:10.1038/s41380-018-0118-1 (2018).

39 Kjaergaard, M., Brander, S. & Poulsen, F. M. Random coil chemical shift for intrinsically disordered proteins: effects of temperature and pH. Journal of biomolecular NMR 49, 139–149, doi:10.1007/s10858-011-9472-x (2011).

40 Kjaergaard, M. & Poulsen, F. M. Sequence correction of random coil chemical shifts: correlation between neighbor correction factors and changes in the Ramachandran distribution. Journal of biomolecular NMR 50, 157–165, doi:10.1007/s10858-011-9508-2 (2011).

41 Schwarzinger, S. et al. Sequence-dependent correction of random coil NMR chemical shifts. Journal of the American Chemical Society 123, 2970–2978, doi:10.1021/ja003760i (2001).

42 Delaglio, F. et al. NMRPipe: a multidimensional spectral processing system based on UNIX pipes. Journal of biomolecular NMR 6, 277–293, doi:10.1007/bf00197809 (1995).

43 Lee, W., Tonelli, M. & Markley, J. L. NMRFAM-SPARKY: enhanced software for biomolecular NMR spectroscopy. Bioinformatics 31, 1325–1327, doi:10.1093/bioinformatics/btu830 (2015).

44 Takegoshi, K., Nakamura, S., & Terao, T. (2001). 13C–1H dipolar-assisted rotational resonance in magic-angle spinning NMR. Chem. Phys. Lett. 344, 631–637, doi:10.1016/S0009-2614(01)00791-6 (2001).

